# Prevention of Drug-Induced Lung Fibrosis via Inhibition of the MRTF/SRF Transcription Pathway

**DOI:** 10.1101/2021.09.09.459118

**Authors:** Kendell M. Pawelec, Megan Varnum, Jack R. Harkema, Bruce Auerbach, Scott D. Larsen, Richard R. Neubig

## Abstract

Drug-induced lung fibrosis is a debilitating disease, linked to high morbidity and mortality. A number of drugs can cause fibrosis, many of which are used to treat cancer, including chemotherapy agents and immune checkpoint inhibitors. The MRTF/SRF transcription pathway has been proposed as a potential therapeutic target, as it is critical for myofibroblast differentiation, a hallmark of fibrosis. In human lung fibroblasts, the MRTF/SRF pathway inhibitor, CCG-257081, effectively decreased mRNA levels of downstream genes: smooth muscle actin and connective tissue growth factor, with IC_50_s of 4 and 15 μM, respectively. The ability of CCG-257081 to prevent inflammation and fibrosis, measured via pulmonary collagen content and histopathology, was tested in a murine model of chemotherapy-induced lung fibrosis. Animals were given intraperitoneal bleomycin for four weeks, and concurrently dosed with CCG-257081 (0, 10, 30, and 100 mg/kg PO), a clinical anti-fibrotic (nintedanib), or clinical standard of care (prednisolone). Mice treated with 100 mg/kg CCG-257081 gained weight vs. vehicle-treated control mice, while those receiving nintedanib and prednisolone lost significant weight. Hydroxyproline content and histological findings in tissue of animals on 100 mg/kg CCG-257081 were not significantly different from naive tissue, indicating successful prevention. Measures of tissue fibrosis were comparable between CCG-257081 and nintedanib, but only the MRTF/SRF inhibitor decreased plasminogen activator inhibitor-1 (PAI-1), a marker linked to fibrosis, in bronchoalveolar lavage fluid. Prednisolone led to marked increases in lung fibrosis. This study demonstrates the potential use of MRTF/SRF inhibitors to prevent drug-induced lung fibrosis in a clinically relevant model of drug-induced disease.

## 1 Introduction

Drug-induced lung fibrosis has been acknowledged as a growing problem that is often not recognized until after FDA-approval for new drugs. Drugs that have been identified to cause lung fibrosis include those that treat cancer, anti-rheumatic drugs (DMARDs), antibiotics, and non-steroidal anti-inflammatory agents (1). The widespread use of these drugs causes long-term morbidity and mortality to patients and can permanently decrease quality of life (2, 3).

The burden of drug-induced lung fibrosis on patients is high and often falls on cancer survivors, as drugs linked to lung fibrosis include chemotherapy agents, such as bleomycin, immune checkpoint inhibitors, etc. Since lung fibrosis is both progressive and debilitating, the average patient life expectancy is four to five years after diagnosis (2). Like all fibrotic diseases, pulmonary fibrosis is caused by an excess of extracellular matrix (ECM) deposition, namely collagen, within the lung. Although initiated by diverse mechanisms, the process generally begins with inflammatory responses (4). These drive the critical transition from a normal healthy lung fibroblast to a myofibroblast – a rapidly dividing cell responsible for increased collagen deposition in the lungs (5). Without the intervention of therapeutics, the build-up of ECM in the lungs impairs tissue elasticity and impedes gas exchange, eventually leading to respiratory failure. While two anti-fibrotics have been clinically approved, they only slow the progression of the disease. The definitive treatment for end-stage lung fibrosis remains lung transplant, which has a median survival as low as 50% at five years post-operation (6). Given the poor prognosis, it is imperative to prevent the development of lung fibrosis before it can progress to end stage disease.

There are currently no clinical therapies approved to prevent or treat drug-induced lung fibrosis. Clinical oncologists treat the symptoms of lung fibrosis with steroids, but these drugs have limited efficacy in the clinic (1). Steroids reduce inflammation, but do not treat the underlying disease and therefore have no effect on measures of fibrosis, and they carry risk of serious side-effects with chronic use. Approved anti-fibrotics, including pirfenidone and nintedanib, are available for other types of pulmonary fibrosis, such as idiopathic pulmonary fibrosis (IPF). However, they are not yet approved for prevention of drug-induced pulmonary fibrosis.

A novel treatment approach for fibrotic diseases targets the transition from fibroblast to myofibroblast, a hallmark of fibrosis through inhibition of the MRTF/SRF transcription pathway. The initiating event for fibrosis can be varied such as reactive oxygen species, DNA damage and inflammation, all of which rely on multiple signaling pathways. Given the diversity of fibrotic pathways and redundancy between pathways, it remains difficult to quiet all of these mechanisms simultaneously. However, Rho signaling through the MRTF/SRF transcriptional switch is a critical common pathway in most fibrotic diseases (5, 7, 8), making it an attractive target for therapeutics. Inhibitors of the MRTF/SRF pathway have demonstrated an ability to halt the progression of fibrosis in a variety of tissues, including lung (9), skin (10-12), and conjunctiva (13).

Most fibrotic diseases are well-established at the time of diagnosis. However, with drug-induced fibrosis, the timing of the initiating insult is known so there is an opportunity for preventive treatment in the clinic. The chemotherapy drug, bleomycin, is well known for causing lung fibrosis during cancer treatment. It has been used in animal models of fibrotic diseases (e.g. IPF and scleroderma), but the pathophysiology of these models, relying on direct administration of drug to tissue, differs greatly from clinical drug-induced lung fibrosis. Alternatively, systemic administration of a pro-fibrotic drug like bleomycin should more closely mimic the inflammatory and fibrotic processes seen in humans in the context of drug-induced fibrosis (14). Taking bleomycin as a model drug, the goal of the current study was to demonstrate the potential of MRTF/SRF inhibitor, CCG-257081, to prevent the development of lung fibrosis in a clinically translatable disease model. Further, the in vivo efficacy of CCG-257081 was compared with steroid treatment, the clinical standard of care, and an anti-fibrotic clinically approved for IPF (nintedanib). An effective and well tolerated preventive therapy for drug-induced lung fibrosis would fill an unmet medical need that could ensure continued quality of life for patients by eliminating the formation of this lifelong and devastating morbidity.

## 2 Materials & Methods

The MRTF/SRF pathway inhibitor, CCG-257081 (12), was supplied by the Vahlteich Medicinal Chemistry Core (University of Michigan, Ann Arbor, MI, USA). For in vitro experiments, it was stored as a 10 mg/ml stock in dimethyl sulfoxide (DMSO, Fisher Scientific, Waltham, MA).

### 2.1 In vitro efficacy: qPCR

Normal human lung fibroblasts (Lonza, Basel, Switzerland) were cultured in DMEM (high glucose, containing 10% heat-inactivated fetal bovine serum (FBS, qualified, USDA-approved regions, Gibco 10-437-010)) and 1% Penicillin-Streptomycin (Gibco 15-140-122); all cells were used before passage 6. To evaluate changes to mRNA expression, 10,000 cells were plated in each well of a 96-well plate for 24 hours before changing the media to DMEM containing 0.5% FBS and 10 ng/ml TGF-β (R&D Systems, Minneapolis, MN, #240-B-002). Concurrently, CCG-257081 (1, 3, 10 μM) or DMSO was added; total DMSO was constant across groups (1%). At 24 hours after addition of TGF-β and CCG-257081, the cells were lysed in iScript RT-qPCR lysis buffer (Biorad, Hercules, CA, cat #170-8898). SYBR Green qPCR was performed using a ViiA7 (A&B Biosystems, Waltham, MA) with iTaq Universal SYBR Green One Step Kit (BioRad, #1725151) following the manufacturer’s instructions. All experiments were repeated three times (n=3 each) and Ct values were analyzed relative to GAPDH expression. Primer sequences used were as follows: *GAPDH*: 5′-GAAGGTGAAGGTCGGAGTCA-3′, 3′CTGGGGAAGTAACTGGAGTT-5′, *CTGF*: 5′-CAGCATGGACGTTCGTCTG-3′, 3′-GGTTCCTGGTTTGGCACCAA-5′, and *ACTA2*: 5′-AGCCCAGCCAAGCACTG-3′, -3′GAGACATTCCGGCCGAAAC-5′.

### 2.2 Preventive model

#### 2.2.1 Animal model

All animal work was conducted by the Michigan State University in Vivo Facility (East Lansing, MI, USA). This study was conducted in accordance with the current guidelines for animal welfare (15). The procedures used were reviewed and approved by the Institutional Animal Care and Use Committee at Michigan State University.

Male mice (C57BL/6J, Charles River Labs) were used at approx. 8.5 weeks of age, as female mice do not develop robust lung fibrosis in bleomycin models (16). Mice were housed individually in solid bottom cages using corncob bedding. All animals were acclimated for 20 days, allowing for stable body weight gain. At the first day of dosing, they weighed between 19.5 - 25.5 g. Throughout the study, animals were maintained on an automated 12/12 hour light/dark cycle. All animals had access to standard rodent chow and fresh water ad libitum.

#### 2.2.2 Formulation

Bleomycin (GoldBio, St. Louis, MO, #B-910-1G) was dissolved in sterile phosphate buffered saline (PBS, Gibco) at 3 mg/ml (15 mg/kg) and sterile filtered. It was stored at -20°C in aliquots appropriately sized for the individual dosing days. The formulation was brought to room temperature just prior to use.

All compounds were prepared weekly and stored at 4°C until just prior to use. CCG-257081 was made up as 2, 6, and 20 mg/mL, corresponding to 10, 30 and 100 mg/kg, in 5% DMSO in polyethylene glycol (PEG)-400. Nintedanib (AstaTech, Bristol, PA, #43021) was prepared in 0.5% methylcellulose (viscosity 4000 cP, MilliporesSigma, St. Louis, MO) at 6 mg/mL (30 mg/kg) and protected from light. Prednisolone (Acros Organics, Geel, Belgium, AC449470010) was prepared in 0.5% methylcellulose. The original formulation was prepared at 3 mg/mL to deliver a dose of 15 mg/kg. However, due to excessive weight loss in animals, on Day 8, the dose was decreased to 1 mg/ml (5 mg/kg) and remained at 1 mg/ml throughout the remainder of the study. The formulations of nintedanib and prednisolone were chosen based on previously published literature in prevention studies (17, 18).

#### 2.2.3 Study Design

Mice were randomly assigned to 6 groups: vehicle (5% DMSO/PEG-400, n=10), 10 mg/kg CCG-257081 (n=15), 30 mg/kg CCG-257081 (n=15), 100 mg/kg CCG-257081 (n=10), 30 mg/kg nintedanib (n=10), and prednisolone (15 mg/kg Days 1-7, 5 mg/kg Days 8-42, n=10). Bleomycin was administered intraperitoneally (IP) at 15 mg/kg twice weekly over 4 weeks to induce lung fibrosis (14). Compounds were administered to mice via once daily oral gavage, starting 24 hours prior to the first dose of bleomycin and continuing over 6 weeks (42 days). Mice were observed daily, and pre-dose clinical observations were recorded. Body weights were collected two times per week.

On Day 42, the final day of dosing, blood (∼250μL) was collected 2 hours post-dose from groups receiving CCG-257081 (6 mice per group) and plasma prepared for quantification of steady state dosing levels. Mice were euthanized on Day 43. Lungs were removed and bronchoalveolar lavage fluid (BALf) was collected. The left lung and BALf were snap frozen in liquid nitrogen and stored at -80°C until analysis. The right lung was inflated and fixed in 10% neutral buffered formalin for subsequent histopathology. Age-matched mice that had not received bleomycin were used as naive controls.

### 2.3 Hydroxyproline content

Snap frozen lung tissue, harvested at Day 43, was diced and added to 200 μl water and 400mg of stainless-steel beads (0.9-2 mm), before homogenizing in a Bullet Blender (Next Advance, Troy, NY, USA) for 4 minutes at speed setting 10. Homogenized tissue was diluted with water (1:10) and added to an equal volume of 37% HCl. The sample was hydrolyzed at 120°C for 3 hours prior to adding 15μl of sample, in duplicate, to a 96-well plate. Hydroxyproline content was measured using a colorimetric test kit (Sigma, MAK008-1KT), per the manufacturer’s instructions, and absorbance was read at 450 nm using a BioTek plate reader. Recorded values are an average of two independent repeats of the assay and are compared to naive lung tissue.

### 2.4 Histopathology

Fixed lung tissue was trimmed and embedded in parafilm prior to sectioning to 10 μm. Tissue sections were histochemically stained with hematoxylin & eosin (H&E) for routine microscopic detection of histopathology and with Masson’s Trichrome to identify areas of fibrosis. Other sections were immunohistochemically stained for lung myofibroblasts containing α-smooth muscle actin (α-SMA; rabbit anti-α-smooth muscle actin, #ab5694, Abcam, Cambridge, MA). Tissue sections were microscopically examined, and lung lesions were semi-quantitatively scored for inflammation, hyperplasia of type 2 alveolar epithelial cells, and fibrosis by a board-certified veterinary pathologist with extensive experience in respiratory pathology of laboratory animals (J.R.H.) and who was without knowledge of individual animal treatment history (“blinded assessment”). Individual lungs were semi-quantitatively graded for lesion severity as % of total pulmonary tissue examined based on the following criteria: (0) no changes compared to control mice; (1) minimal (<10%); (2) slight (10–25%); (3) moderate (26–50%); (4) severe (51–75%); or (5) very severe (>75%) of total area affected.

### 2.5 PAI-1 ELISA

BALf was assayed for plasminogen activator inhibitor-1 (PAI-1) concentration using an ELISA kit (PAI-1 (*SERPINE1*) Mouse ELISA, Thermo Fisher, EMSERPINE1). The assay was conducted according the manufacturer’s instructions. All samples were diluted 1:15 and run in duplicate. BALF from 6-8 individuals per group, chosen at random, were quantified.

### 2.6 Statistics

All statistical analysis was done using Graphpad Prism, version 9.1.0. In all cases, a p value <0.05 was considered statistically significant. Unless specified, significance was tested via ANOVA, with a Dunnett’s post-test for pairwise comparisons between groups. For body weight data, mixed effects analysis was performed to account for missing data due to animal loss over the study. Histopathology was assessed via a non-parametric Kruskal-Wallis test.

## 3 Results

### 3.1 Inhibition of MRTF/SRF-Regulated Transcription in Lung Fibroblasts

MRTF/SRF pathway inhibitors have previously been shown to reduce myofibroblast differentiation in skin fibroblasts (10, 11). To validate the same response in lung, normal human lung fibroblasts were stimulated with pro-fibrotic TGF-β in the presence of a known MRTF/SRF pathway inhibitor (CCG-257081) in concentrations ranging from 1-10 μM. Levels of mRNA for two genes under direct transcriptional control by MRTF/SRF, α-smooth muscle actin (*ACTA2*) and connective tissue growth factor (*CTGF*) (19) were evaluated (Figure 1A). CCG-257081 reduced mRNA for both genes in a concentration-dependent manner (Figure 1B). The IC_50_ values in human lung fibroblasts were approximately 4 and 15 μM for *ACTA2* and *CTGF*, respectively. The reduction in ACTA2 at 10 mM CCG-257081 was significant (p < 0.001).

**Figure 1:**
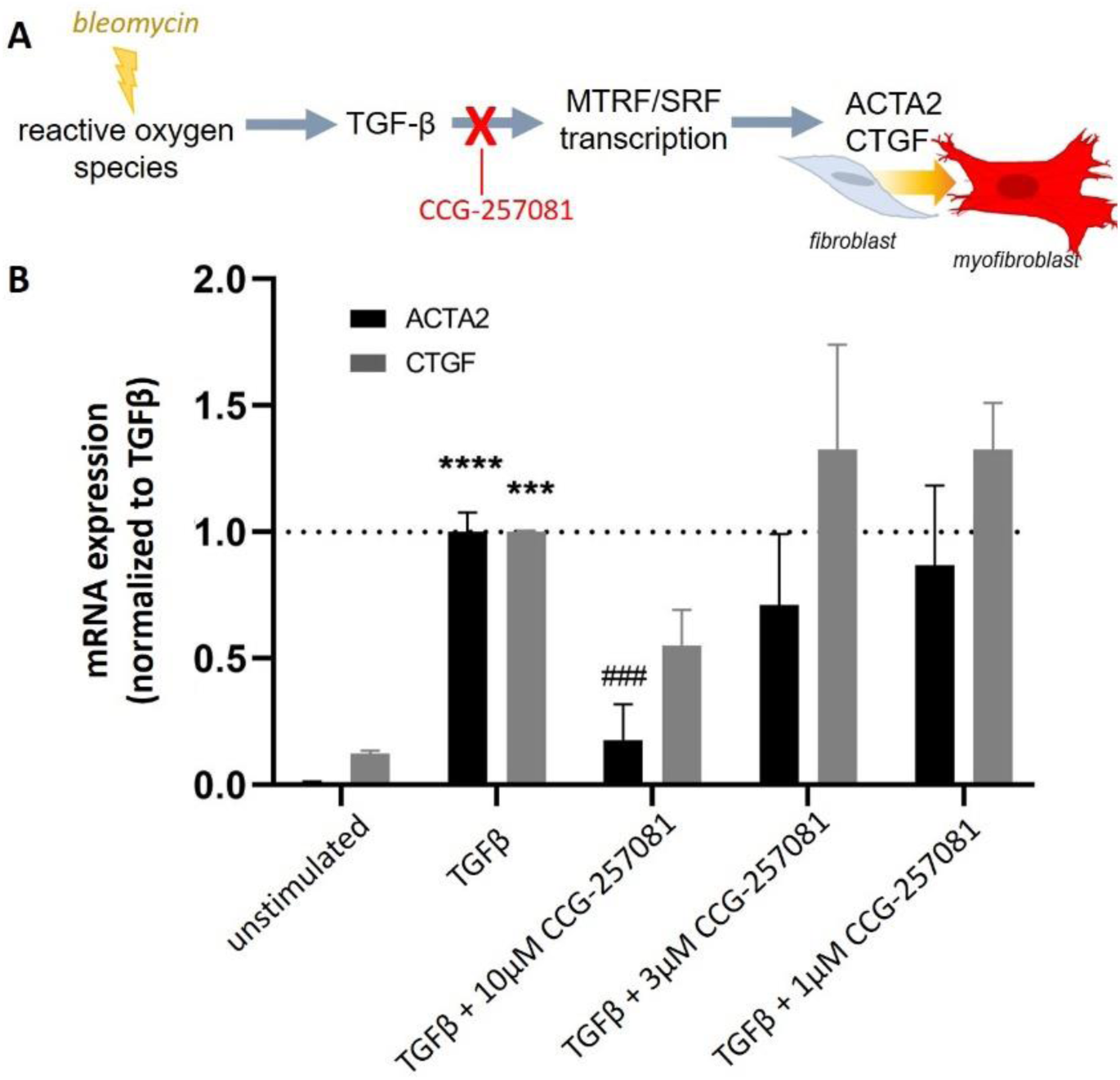
Inhibition of MRTF/SRF-regulated transcription in human lung fibroblasts reduces mRNA levels of genes related to myofibroblast differentiation. **A** Schematic of the role MRTF/SRF inhibitors play during in vivo drug-induced fibrosis and **B** the reduction in mRNA expression of actin (*ACTA2*) and connective tissue growth factor (*CTGF*) with CCG-257081 administration after treatment with pro-fibrotic stimulus, TGF-β. **** p < 0.0001, *** p < 0.001 vs unstimulated; ### p < 0.001 vs TGFβ-stimulated.

### 3.2 Prevention of Drug-induced Lung Fibrosis

Having validated that the MRTF/SRF inhibitor was able to prevent the myofibroblast transition in vitro, the hypothesis was tested in vivo in a murine model designed to mimic clinical events leading to drug-induced lung fibrosis. Namely, the pro-fibrotic stimulus (bleomycin sulfate) was administered systemically via intraperitoneal injection over 4 weeks. Body weight was monitored during the study (Figure 2). In the more commonly used model of lung fibrosis, where bleomycin is administered by intratracheal instillation, mice show rapid and substantial loss of body weight (9). All animals received bleomycin treatment. However, with the milder lung injury induced by this dose of parenteral bleomycin, animals treated with bleomycin and vehicle ((5% DMSO/PEG400) only had no net change in weight over the 6-week period. Animals which did not receive any bleomycin treatment would be expected to have an increase in body weight of 25-30% over this time frame. Prednisolone, the clinical standard of care, caused a substantial and rapid loss of weight over the first week, necessitating a reduction in dose level from 15 to 5 mg/kg. Despite the dose reduction, animals did not recover body mass, even after bleomycin treatment was stopped at day 28. The group treated with nintedanib, an approved anti-fibrotic drug, also experienced substantial weight loss in the first week, but returned to baseline levels by the end of the study. The animals treated with the highest dose of CCG-257081 (100 mg/kg) did not lose any weight and experienced a significant gain in weight above the bleomycin + vehicle control. The plasma concentrations of CCG-257081 in mice at 2 hours after the final dose of compound were 2.98 ± 1.2 μM, 12.2 ± 4.4 μM, and 28.3 ± 4.9 μM for 10, 30 and 100 mg/kg dosing, respectively.

**Figure 2:**
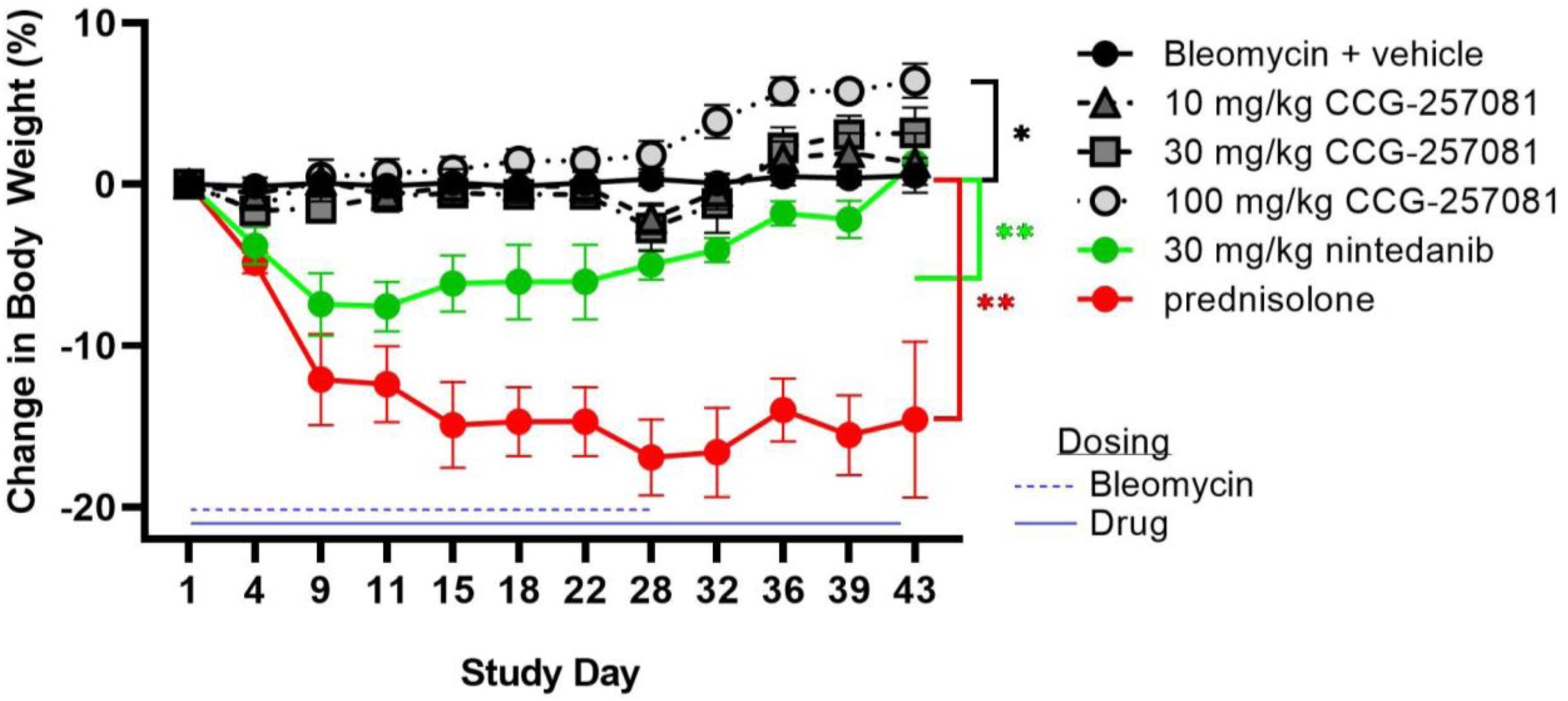
Change in body weight during drug treatments. All animals received bleomycin treatment from day 1 -28 and test agents from days 1-42. Stress on the animals was observed in the rapid weight loss during the first week in the nintedanib and prednisolone groups, and a lack of significant weight gain over the 6 week study in all groups except 100 mg/kg CCG-257081. Due to excessive weight loss, the dose of prednisolone was lowered from 15 mg/kg to 5 mg/kg. Weight change time courses, which are significantly different from the vehicle control, are marked (*p < 0.05, **p < 0.01).

At 6 weeks, 2 weeks after cessation of bleomycin dosing, lung tissue was microscopically assessed for drug-induced pulmonary histopathology by a board-certified veterinary pathologist (J.R.H.) who was blinded to the treatment given. Pulmonary lesions were restricted to perivascular and subpleural regions of the lung. The lesions were characterized by chronic alveolitis composed of interstitial and alveolar fibrosis and an associated mononuclear inflammatory cell infiltrate consisting of monocytes, lymphocytes, and hypertrophic vacuolated alveolar macrophages, and areas of hyperplasia/hypertrophy of type 2 alveolar epithelial cells (AT2). No such lung lesions were present in the lungs of control mice. Animals treated with prednisolone had the most severe bleomycin-induced lung lesions with severity scores significantly greater than other experimental groups, i.e., a median score of 4 for inflammation (Figure 3A) and AT2 hyperplasia (Figure 3B). In contrast, there was a was a dose-dependent decrease in histopathology with CCG-25708 treatment, with minimal to no bleomycin-induced lung lesions in mice receiving the highest dose, 100 mg/kg.

**Figure 3:**
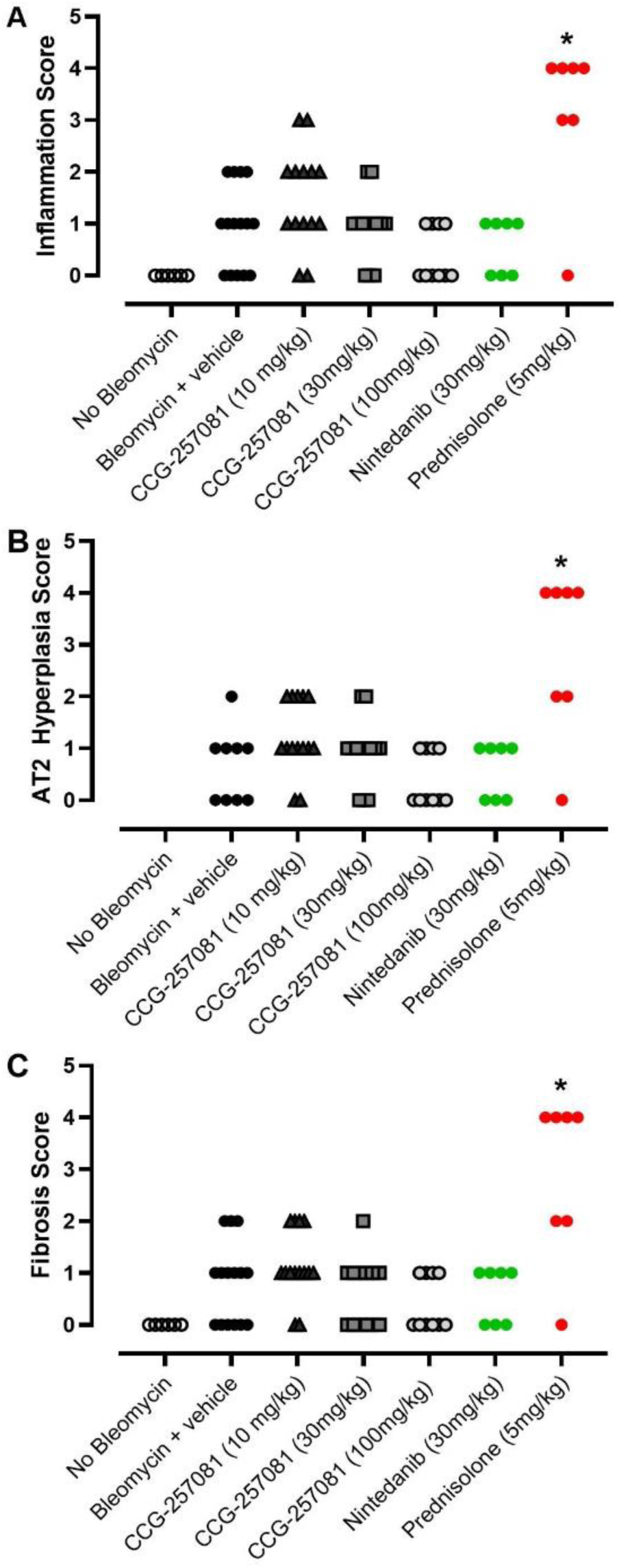
CCG-257081 reduces inflammatory responses inherent to bleomycin-induced fibrotic disease. Histopathology scoring of **A** inflammation, **B** AT2 Hyperplasia and **C** fibrosis showed a trend in lower scores with higher concentrations of CCG-257081. *Note: no data was recorded for the “no bleomycin” control group in AT2 Hyperplasia*. The severity scores were: 0, no significant findings; 1, minimal; 2, mild; 3, moderate; 4, marked; 5, severe. * p < 0.05.

Fibrosis in lung tissue was assessed by histopathology/semi-quantitative severity scores and quantified by collagen/hydroxyproline content in the lungs. Fibrosis scores (Figure 3C), mirrored the inflammation scores, with prednisolone-treated mice demonstrating the highest levels of fibrosis (median: 4). Similar to the inflammation severity scores, the median severity score for fibrosis in mice treated with CCG-257081 at 100 mg/kg was 0. The fibrosis observed was interstitial and alveolar, with varying degrees of fibrotic obscuring of alveolar air spaces. Staining of the lung tissue confirmed the high collagen content within fibrotic lesions in the lung (Figure 4) and that myofibroblasts were present in the lesions (Supplemental Fig. 2).

**Figure 4:**
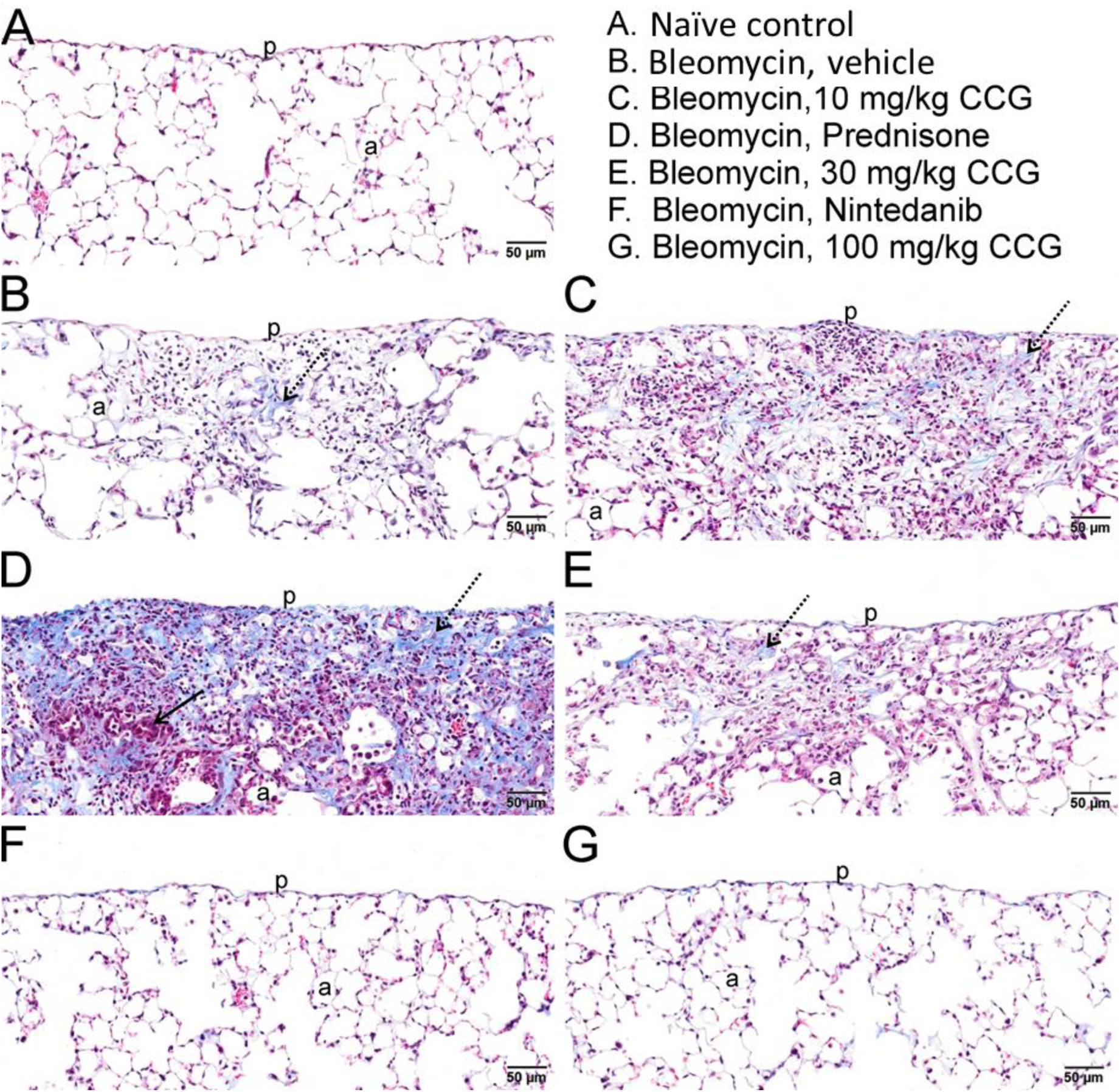
Light photomicrographs of lung tissue from mice in experimental groups: **A** Naïve control group, **B** Bleomycin + vehicle, **C** Bleomycin with low dose CCG-257081 (10 mg/kg), **D** Bleomycin with Prednisone treatment, **E** Bleomycin with medium dose CCG-257081 (30 mg/kg), **F** Bleomycin with Nintedanib treatment, and **G** Bleomycin with high dose CCG-257081 (100 mg/kg). Tissue sections histochemically stained with Masson’s Trichrome (blue: collagen, red: red blood cells, gray: alveolar space). Bleomycin-induced subpleural chronic alveolitis with increased collagen deposition (fibrosis; blue chromogen/stippled arrows) was observed, that was most severe in D (Prednisone treatment). No subpleural fibrotic lesions were noted with high dose CCG-257081 (G) or Nintedanib (F). p, pleural surface of lung; a, alveolar parenchyma; solid arrow, alveolar type II epithelial hyperplasia/hypertrophy.

Collagen deposition in the lungs, measured via hydroxyproline content, confirmed the histology results (Figure 5A). Prednisolone-treated animals had significantly more collagen deposition than naive animals. Unfortunately, the relatively modest injury observed with the parenteral bleomycin and a large amount of scatter in the bleomycin + vehicle control group limited the conclusions which could be drawn. However, there was a significant difference in the reduction of collagen content from low to high CCG-257081 doses and between prednisolone and CCG-257081 (100 mg/kg). There was also no statistical difference between collagen levels in naive tissue and CCG-257081 at 100mg/kg.

**Figure 5:**
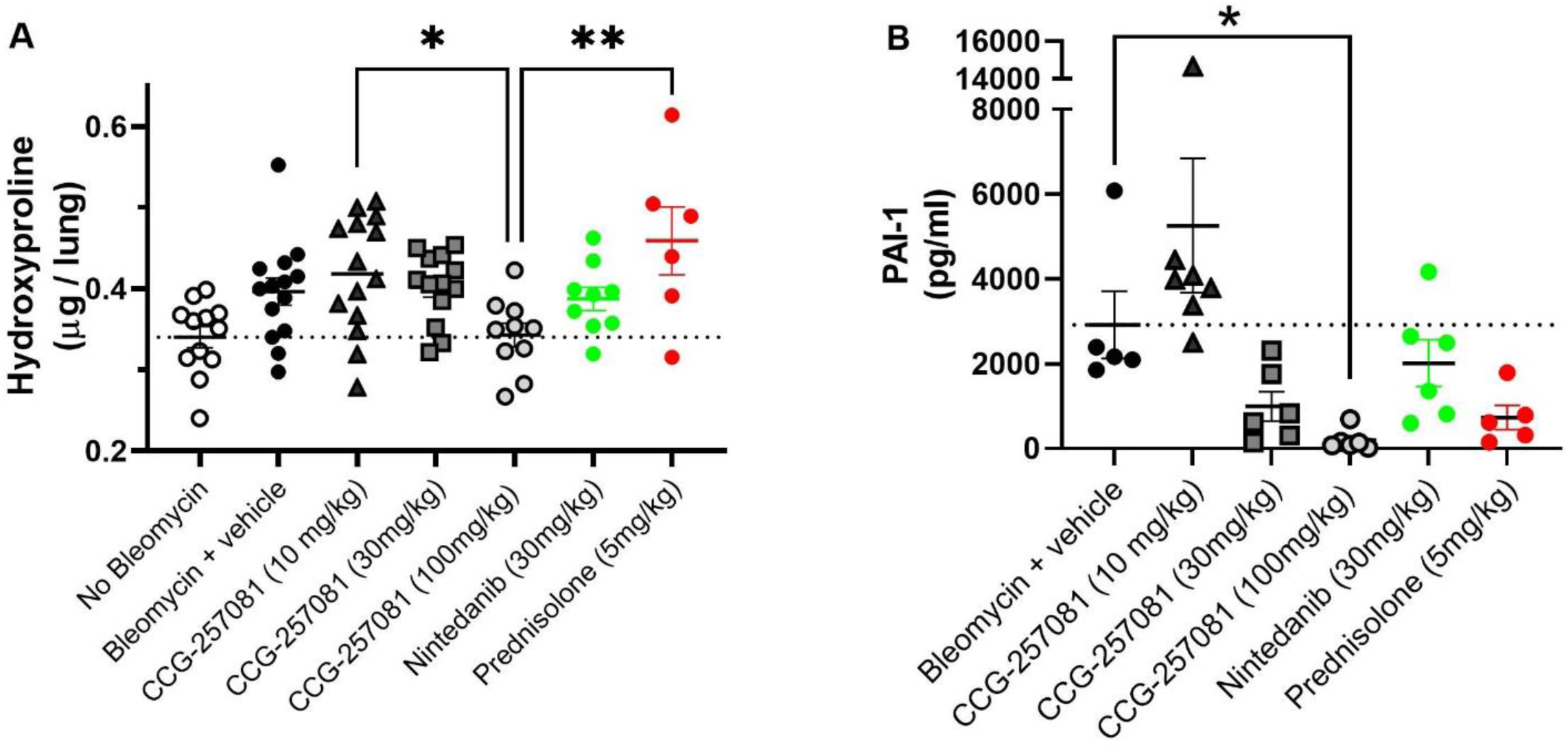
Fibrotic markers were significantly decreased by MRTF/SRF inhibitors. **A** At 100 mg/kg CCG-257081, collagen content in the lungs was not significantly different from naive tissue (no bleomycin), while animals with prednisolone treatment had significantly greater amounts of collagen. **B** The pro-fibrotic biomarker, PAI-1, was significantly reduced in BALf by treatment with CCG-257081 at 100 mg/kg, but was not significantly reduced by any other treatment, including nintedanib or prednisolone. *p < 0.05, **p < 0.01.

BALf was also assayed for the pro-fibrotic mediator, plasminogen activator inhibitor-1 (PAI-1). Transcription of the gene that encodes for PAI-1 (SERPINE1) is directly controlled by MRTF/SRF (19). PAI-1 is also a known marker and potential drug target associated with inflammation and fibrotic disease (20). CCG-257081 significantly reduced PAI-1 in BALF, with a greater than 15-fold reduction at 100 mg/kg compared to vehicle control. In contrast to collagen content, PAI-1 levels tended to be lower in animals treated with prednisolone compared to vehicle alone, but nintedanib dosing had no significant effects on PAI-1 (Figure 5B).

## 4 Discussion

Prevention of drug-induced lung fibrosis is a significant unmet medical need that is growing in importance, particularly in the treatment of cancer, but preventive therapies must be well-tolerated. Agents implicated in the disease include bleomycin, gemcitabine, epidermal growth factor receptor kinase inhibitors, and immune checkpoint inhibitors (1). The severe burden to patients diagnosed with drug-induced lung fibrosis and the poor prognosis of end-stage disease make it imperative that treatment options be identified. Drug-induced lung fibrosis is difficult to diagnose as the clinical and histopathology patterns vary with drugs and with the individual (1). However, the chemotherapy drug, bleomycin sulfate, results in the highest incidence of drug-induced interstitial lung disease (1) and has been extensively studied as a fibrosis-inducing agent in animal models (14, 21). This is why it was chosen as the representative drug model for this study.

Hallmarks of bleomycin-induced lung fibrosis in the clinic include patchy inflammation, reactive epithelial hyperplasia, infiltration of inflammatory cells and remodeling of the extracellular matrix (ECM) (21). Ideally, animal models that are used assess novel treatment therapies would recapitulate these features. However, the majority of in vivo models of bleomycin-induced fibrosis use direct intratracheal administration of drug to the lung or injection into the skin, eliciting a massive local fibrotic response that is not clinically relevant. The current study sought to use a more translatable model for inducing lung fibrosis by administering bleomycin systemically. Accordingly, changes in body weight during the trial and the amount of fibrotic tissue formed (see Figures 2 and 4) were relatively subtle (< 20%) compared to other commonly used models. In contrast to studies with intratracheal bleomycin, which cause substantial and rapid losses of body weight (9), we saw only prevention of weight gain by the animals, rather than a true weight reduction. Evaluation of the histopathology revealed subpleural lesions that mimicked the clinical pathology of drug-induced interstitial lung disease far better than do those induced by intratracheal instillation (21). Unlike in mice, in humans, when inflammatory responses lead to damage of the lung tissue structure, the resulting fibrosis is permanent (22). Thus, any fibrotic scarring in the lungs, even modest changes, can contribute to a lifelong decrease in lung function and impairment.

The clinical standard of care when drug-induced lung fibrosis is identified is to stop drug administration if diagnosed during treatment. This is often followed by administration of steroids, as they are known to decrease inflammatory responses that may up-regulate fibrotic pathways. However, the use of steroids for treatment of fibrosis remains controversial. The effects in animal models of lung fibrosis are dependent on dose and timing of administration, with high doses at early time points leading to increased infiltration of lungs by inflammatory cells (23). Clinical trial results have also been inconsistent, demonstrating, at best, very limited benefits of steroids in drug-induced lung fibrosis (1). In some cases, patients dosed with combination therapies for fibrotic diseases even have increased likelihood of mortality and hospitalization (24).

Despite this situation, the lack of approved therapies for treating drug-induced lung fibrosis still drives the use of steroids in the clinic. For this reason, the current study included a steroid control and the clinical anti-fibrotic drug, nintedanib, although it is not yet approved for this indication. A dose of 15 mg/kg prednisolone was chosen, as prevention studies in literature have reported it safe and effective at eliminating inflammatory responses in rodent models of lung fibrosis induced by bleomycin (17, 25). In particular, the action of corticosteroids was differentiated from anti-fibrotics by working only on inflammation, decreasing inflammatory cytokines and pulmonary edema, with no effect on cellular infiltration or pulmonary fibrosis (17).

In the present study, administration of prednisolone resulted in severe adverse events early on, consistent with catabolic effects on skeletal muscle. Even after decreasing the dose to 5 mg/kg, animals did not regain weight, and they had increased levels of inflammation, lung fibrosis and hydroxyproline content. A link has been established between high collagen content in the lungs and high levels of PAI-1, a marker associated with both inflammatory and pro-fibrotic pathways (20, 26). However, PAI-1 was decreased by prednisolone. Since multiple cell types can produce PAI-1, including alveolar epithelial cells and macrophage populations, prednisolone may quiet PAI-1 expression in a subset of these cells, but not enough to inhibit a fibrotic response (20). Given the known sensitivity of steroid efficacy to dose and timing (23), it is likely that differences in the animal model, particularly the dose and route of bleomycin administration, played a role in the increased fibrosis over vehicle controls. Regardless, there is clearly a need for alternative therapies for drug-induced lung fibrosis, such as MRTF/SRF inhibitors.

### 4.1 Inhibition of MRTF/SRF across tissues

The CCG series of MRTF/SRF transcription pathway inhibitors down-regulate mRNA transcription of target genes in a variety of tissues (10, 11, 13). In the present study, the direct action of the MRTF/SRF pathway inhibitor, CCG-257081, was confirmed in lung fibroblasts. Two genes critical to myofibroblast differentiation during fibrosis are those for αSMA (*ACTA2*) and connective tissue growth factor (*CTGF*). An in vitro concentration response curve showed that both of these MRTF/SRF-regulate genes are reduced by CCG-257081 in normal human pulmonary fibroblasts, confirming the ability of MRTF/SRF inhibition to control the fibrotic switch in lung.

*ACTA2* was strongly inhibited (IC_50_ ∼ 4 µM), while *CTGF* inhibition had an IC_50_ of ∼15 µM. Notably, plasma concentrations of CCG-257081 at 2 hours after dosing were 12 and 28 μM for the two higher doses of compound used in this study. *ACTA2* (αSMA) transcription is induced late after TGF-β stimulation and can be wholly blocked by ROCK or MRTF/SRF inhibitors (27). In contrast, *CTGF* regulation is more complex, as it is induced early after TGF-β stimulation (within 3 hours), but its transcription is regulated by both MRTF/SRF and SMAD pathways, which are not affected by MRTF/SRF (27). This may explain why CCG-257081 was not as effective at blocking *CTGF* transcription. The clear inhibition of genes downstream of MRTF/SRF in lung fibroblasts, as previously seen in skin fibroblasts, supports the hypothesis that the MRTF/SRF transcriptional switch is critical in pro-fibrotic pathways across multiple tissues.

Recently, the redox-regulated nuclear transcription factor, pirin, was identified as a potential direct target of the CCG series of MRTF/SRF pathway inhibitors (10). In addition to its connection with the pro-fibrotic MRTF/SRF pathway, pirin has also been implicated in modulation of NFκB- (28) and HSF1- (29) regulated gene transcription. These pathways raise the possibility that interaction of CCG compounds with pirin may suppress fibrosis through inflammatory pathways, as well as MRTF/SRF, and potentially contribute to the suppression of inflammation-related readouts in this model. Defining interactions of pirin in fibrotic disease will be important questions for future study.

### 4.2 Role of MRTF/SRF in Drug-Induced Lung Fibrosis

With the goal of identifying novel therapeutics to treat drug-induced lung fibrosis, MRTF/SRF inhibitors were compared to clinically approved anti-fibrotics in a model of systemic bleomycin-induced lung fibrosis. The range of doses for CCG-257081 (10 - 100 mg/kg CCG-257081) was chosen, based on an efficacious dose of 50 mg/kg in treatment models of skin fibrosis (10, 12). At 10 mg/kg, there was no observed efficacy on measures of fibrosis in lung tissue. Further investigation revealed that the plasma levels in mice at this dose were below the in vitro IC_50_ values for *ACTA* and *CTGF*. Plasma levels increased with dose, which corresponded to a reduction in inflammation and fibrosis markers. At 100 mg/kg CCG-257081, there was no significant change in hydroxyproline content or histopathology compared to naive lung tissue, indicating effective protection of the lung tissue. Importantly, animals in this treatment group also had a significant increase in body weight over the vehicle control mice (Figure 2). In contrast, both nintedanib and prednisolone resulted in significant weight loss compared to the mice treated with vehicle only.

As noted, the bleomycin + vehicle group had a modest increase and a high amount of scatter in lung hydroxyproline and histological results, which made it difficult to draw firm conclusions on the effects of bleomycin alone on lung tissue. The low levels of fibrosis induced by systemic administration of bleomycin were expected to produce a more subtle effect than direct application of the bleomycin, but treatment with vehicle can also affect tissue physiology via increased animal handling or increased hydration of the animals. Another variable may be the vehicle chosen for the MRTF/SRF inhibitors, which contained low levels of DMSO. It is known that DMSO has anti-inflammatory properties and that high levels can prevent tissue fibrosis in animal models (30), but the low levels (0.5 mg/kg) administered in the vehicle would be expected to have little effect on the results. Animals dosed with bleomycin and vehicle alone had no change in body weight over the course of the study, far below the expected 20-30% change in weight for naive animals, so there was clear evidence of bleomycin toxicity, albeit milder than is often seen in other studies.

In addition to comparing MRTF/SRF inhibitors to the current clinical standard of care, CCG-257081 was compared to a clinically approved anti-fibrotic drug, nintedanib, not known to act on the MRTF/SRF pathway. As a tyrosine kinase inhibitor, nintedanib disrupts both fibrotic and angiogenic pathways (31, 32). During the current study, animals were given nintedanib at the lowest published dose shown to prevent lung fibrosis (30 mg/kg) (18) to avoid the known gastrointestinal toxicities of the drug (31). Once-a-day dosing of CCG-257081 at 100 mg/kg had a comparable or perhaps slightly better effect vs. nintedanib in qualitative and quantitative measures of fibrosis. Interestingly, the pro-fibrotic marker, PAI-1, was inhibited only by CCG-257081. While nintedanib has anti-inflammatory effects, it does not inhibit cellular infiltration into lungs or decrease the macrophage count in BALf (18). Since macrophages have been identified as a source of PAI-1 secretion in lung fibrosis (20), this may be the reason for no significant effect of nintedanib on PAI-1 compared to the vehicle control. PAI-1 is under the control of MRTF/SRF transcription, likely contributing to the significant decrease in PAI-1 at the higher doses of CCG-257081.

## 5 Conclusion

There is a clear need for well-tolerated preventive treatments for drug-induced lung fibrosis, particularly as the current clinical standard of care, steroid administration, gives very little benefit to patients. MRTF/SRF inhibitors act on a transcriptional pathway central to myofibroblast differentiation, a hallmark of fibrotic disease; a mechanism that plays a role in fibrosis in multiple tissues. In human lung fibroblasts, the MRTF/SRF inhibitor CCG-257081 was able to decrease mRNA expression of both αSMA and CTGF in a concentration-dependent manner. Using a model of modest bleomycin-induced lung fibrosis, designed to replicate the injury seen during systemic chemotherapy treatment, the highest dose of the MRTF/SRF inhibitor CCG-257081 reversed the bleomycin-induced body weight loss, and lung tissue was not significantly different from naive lung tissue in measures of inflammation, collagen content and histopathology scoring. This was significantly better than prednisolone, the clinical standard of care, and comparable to a clinically approved anti-fibrotic agent (nintedanib). The MRTF/SRF inhibitor also strongly suppressed PAI-1 in bronchoalveolar lavage fluid, suggesting that PAI-1 may be a suitable biomarker for tracking drug action. Together, these data support the use of MRTF/SRF inhibitors as potential clinical treatments for preventing drug-induced lung fibrosis, especially in light of the improved tolerability compared to clinical alternatives.

## 6 Acknowledgements

This grant was funded by the National Cancer Institute of NIH (R43CA235823-01). The authors thank Dr. Teresa Krieger-Burke and the MSU In Vivo Facility for assistance with the design and execution of this study.

## Notes

### Competing Interest Statement

KMP, MV, SDL and RRN hold equity in FibrosIX Inc and hold leadership positions in the company. KMP and MV have been/are employees of FibrosIX Inc. and RRN has been paid as a consultant by the company.

