## Supplemental for "Prevention of Drug-Induced Lung Fibrosis via Inhibition of the MRTF/SRF Transcription Pathway"

### 8 Supplemental Information

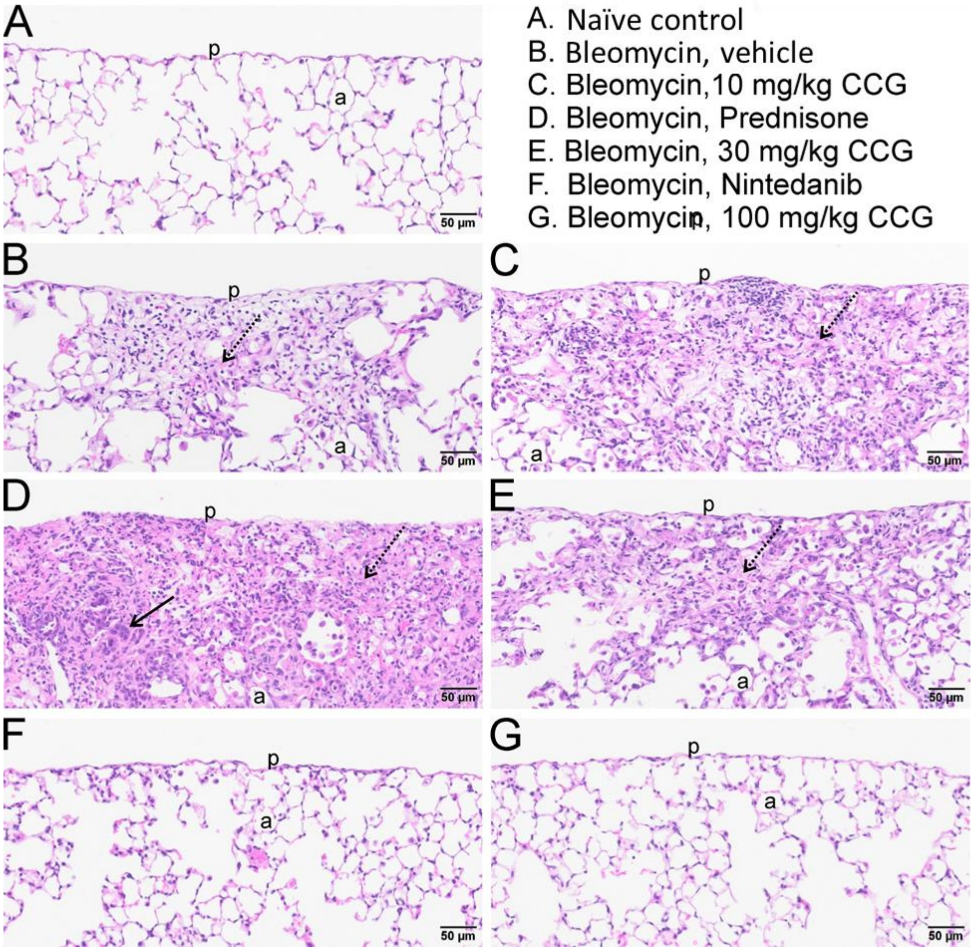


**Supplemental Figure 1:** Lung tissue was affected by treatment, as seen in H&E staining. Representative images of lung tissue (purple: nuclei, pink: extracellular matrix and cytoplasm). Compared to **A** naïve lung tissue, **B** bleomycin+vehicle groups had lesions. Lesions in groups with bleomycin+ CCG-257081, were reduced as the dose increased from **C** 10mg/kg, **E** 30 mg/kg and **G** 100mg/kg, which was comparable to **F** nintedanib. **D** The prednisolone group exhibited the greatest level of fibrosis across all groups. Scale bar is 50 µm. p, Pleura; a, Alveolus; stippled arrow: collagen/fibrosis; solid arrow: epithelial hyperplasia/hypertrophy.

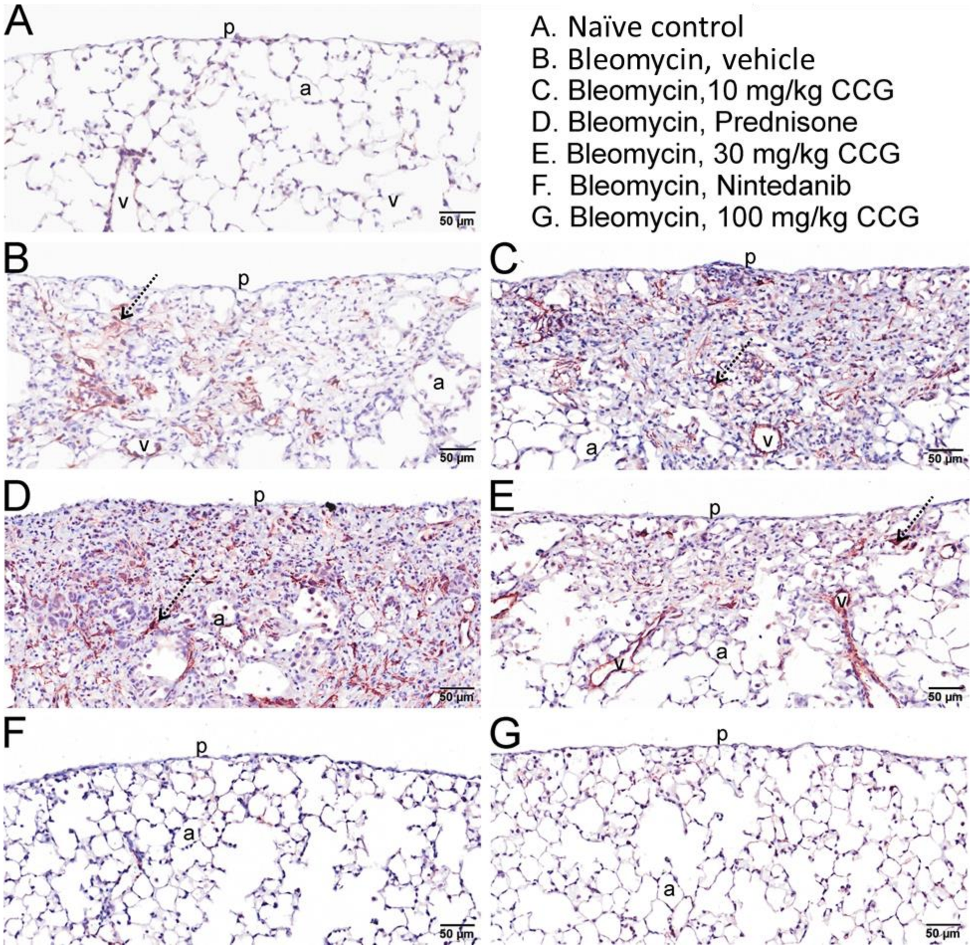


**Supplemental Figure 2:** CCG-257081 and nintedinib reduced myofibroblasts, as measured by αSMA positivity. Representative images of lung tissue (blue: nuclei, brown: αSMA). Compared to **A** naïve lung tissue, **B** bleomycin+vehicle groups had lesions. Lesions in groups with bleomycin+ CCG-257081, were reduced as the dose increased from **C** 10mg/kg, **E** 30 mg/kg and **G** 100mg/kg, which was comparable to **F** nintedanib. **D** The prednisolone group exhibited the greatest level of αSMA staining across all groups. Scale bar is 50 µm. p, Pleura; a, Alveolus; v, blood vessel; stippled arrow: myofibroblast.
